# Electronic Health Record and Genome-wide Genetic Data in Generation Scotland Participants

**DOI:** 10.1101/154609

**Authors:** Shona M. Kerr, Archie Campbell, Jonathan Marten, Veronique Vitart, Andrew McIntosh, David J. Porteous, Caroline Hayward

## Abstract

This paper provides the first detailed demonstration of the research value of the Electronic Health Record (EHR) linked to research data in Generation Scotland Scottish Family Health Study (GS:SFHS) participants, together with how to access this data. The structured, coded variables in the routine biochemistry, prescribing and morbidity records in particular represent highly valuable phenotypic data for a genomics research resource. Access to a wealth of other specialized datasets including cancer, mental health and maternity inpatient information is also possible through the same straightforward and transparent application process. The Electronic Health Record linked dataset is a key component of GS:SFHS, a biobank conceived in 1999 for the purpose of studying the genetics of health areas of current and projected public health importance. Over 24,000 adults were recruited from 2006 to 2011, with broad and enduring written informed consent for biomedical research. Consent was obtained from 23,603 participants for GS:SFHS study data to be linked to their Scottish National Health Service (NHS) records, using their Community Health Index (CHI) number. This identifying number is used for NHS Scotland procedures (registrations, attendances, samples, prescribing and investigations) and allows healthcare records for individuals to be linked across time and location. Here, we describe the NHS EHR dataset on the sub-cohort of 20,032 GS:SFHS participants with consent and mechanism for record linkage plus extensive genetic data. Together with existing study phenotypes, including family history and environmental exposures such as smoking, the EHR is a rich resource of real world data that can be used in research to characterise the health trajectory of participants, available at low cost and a high degree of timeliness, matched to DNA, urine and serum samples and genome-wide genetic information.

## Introduction

GS:SFHS is a large, family-based, intensively-phenotyped cohort of volunteers from the general population across Scotland, UK (www.generationscotland.org). The median age at recruitment was 47 for males and 48 for females, and the cohort has 99% white ethnicity.^1^ The numbers of participants with full phenotype data, genome-wide genotype, consent and mechanism for health record linkage are shown in Figure 1. Research data (baseline and derived subsequent to recruitment) are managed by Generation Scotland in a study database and routine EHR data stored by NHS Scotland in a national database. To create the EHR research dataset, data on GS:SFHS participants was extracted by the NHS National Services Scotland electronic Data Research and Innovation Service (eDRIS), using the CHI number for linkage, then de-identified with a new ID. For research purposes this data is housed in a secure data centre, such as a safe haven. It can be analysed through secure protocols, then deidentified individual-level study and routine medical data brought together for analysis in specific approved research projects (Figure 1). This mechanism is designed to respect the wishes and expectations of the volunteer participants.^2^

**Figure 1.**
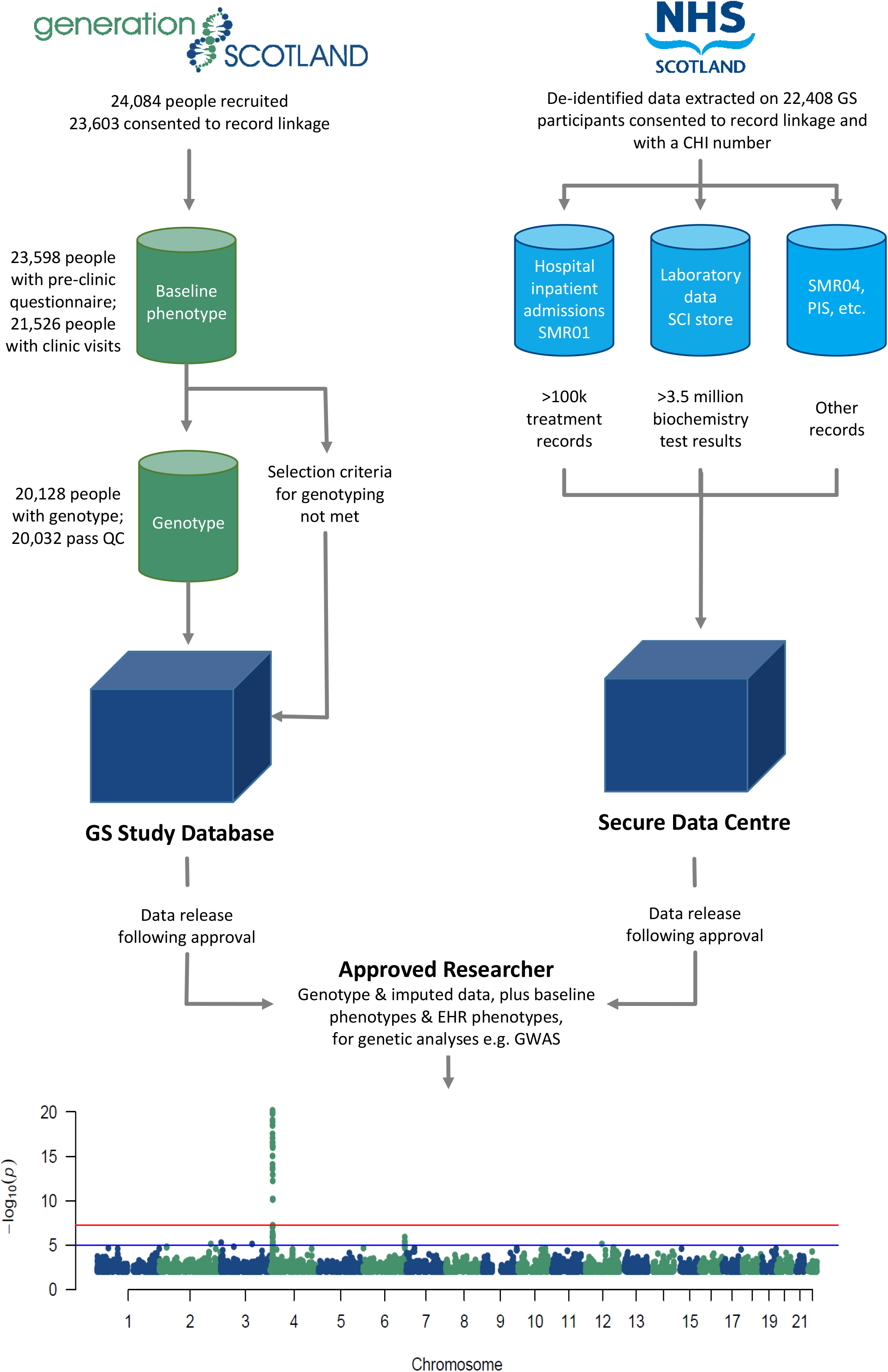
Schematic illustrating the mechanism for research data analyses. The datasets available in Generation Scotland and EHRs are indicated, with numbers of participants and records. The Manhattan plot displays the results of a genome-wide association analysis (GWAS) using genotyped SNPs and EHR-derived serum urate measurements, as an example of how the datasets can be used together in genetic research by approved researchers. The single highest serum urate reading was taken for each participant, with covariates and methods for accounting for relatedness as previously described.^5^ The-log10 (P-value) is plotted on the y-axis, and chromosomal location is plotted on the x-axis. The genome-wide significance threshold accounting for multiple testing (p-value < 5 × 10^-8^) is indicated by a red line, while suggestive significance (p-value < 10^-5^) is indicated by a blue line.

The genome-wide genetic data in GS:SFHS has been used in a large number of research projects across a wide range of study phenotypes that have generated over 100 publications (www.ed.ac.uk/generation-scotland/news-events/publications). Three papers have to date been published on research that used record linkage in GS:SFHS. The first was on the impact of parental diabetes on offspring health, in 2015.^3^ The second (and first example of NHS record linkage for genetic research in GS:SFHS) involved the identification of over 200 cases with atrial fibrillation and matched controls by linkage to hospital episode (Scottish Morbidity Record, SMR01) data (based on International Classification of Diseases (ICD-10) codes), as part of the AFGen Consortium.^4^ The third used linkage to NHS biochemistry data in a genome-wide association study of both directly measured and imputed genotypes.^5^

Extensive and detailed phenotyping, including longitudinal biochemistry data, is of considerable utility in understanding underlying biological or disease mechanisms. However, this data can be difficult and expensive to obtain directly, as laboratory measures require different assays and the quantity of donated samples (e.g. serum, plasma) is finite in a biobank such as GS:SFHS. Collecting longitudinal samples and data in a research setting requires ongoing recontact with study participants, which is both costly and time-consuming. Access to routine EHR data is therefore of great value in genetic research across broad-based medical specialities,^6^ as exemplified by the Electronic Medical Records and Genomics (eMERGE) network^7^ in the USA, and the UK Biobank.^8^

## Materials and Methods

GS:SFHS probands were first approached through their General Practitioner (GP) using the Community Health Index (CHI) database, which has a unique number for each individual in the >96% of the Scottish population registered with a GP.^9^ Those who indicated that they and one or more of their relatives were considering participation were sent an information leaflet, a consent form and a questionnaire. The details of this recruitment of 24,084 participants to GS:SFHS have been described previously.^1^ DNA was extracted from the blood or saliva of participants^10^, and samples were genotyped using a genome-wide SNP array (Illumina OmniExpressExome).^5^ A subset of participants was selected for genotyping, consisting of those individuals who were born in the UK, had Caucasian ethnicity, had full baseline phenotype data available from a visit to a GS:SFHS research clinic in Aberdeen, Dundee, Glasgow or Perth, and had consented for their data to be linked to their NHS records.^5,11^ The total number of participants genotyped was 20,128, of which 20,032 passed additional genetic quality control filtering.

Scotland has some of the most comprehensive health service data in the world. Few other countries can lay claim to national indexed data of such high quality and consistency. Timelines for individual administrative datasets useful in research are given at www.adls.ac.uk/wp-content/uploads/ISD-timelines.pdf, with (for example) General Acute / Inpatient Scottish Morbidity Record (SMR01) data available from 1981. Coverage dates for biochemistry EHRs vary regionally across Scotland due to different dates of implementing storage of records in electronic format across the NHS Area Health Boards. Records in NHS Greater Glasgow and Clyde are available from May 2006 onwards, while some NHS Tayside records (covering the Dundee and Perth GS:SFHS recruitment areas) go back as far as 1988. Data can therefore be accessed for most participants from well before the period of recruitment to the GS:SFHS cohort (2006-2011) and subsequent to participation in the study, up to within a few months of the date of a data release. The resource includes contemporary measures that reflect current tests and treatments. Longitudinal research is therefore feasible, with mechanisms also in place for re-contact of GS participants for targeted follow-up,^11^ including recall-by-genotype studies,^12^ enabling detailed research on chronic conditions and long-term outcomes.

## Dataset Validation

The Generation Scotland phenotype, genotype and imputed data have been subject to extensive quality control and are research ready. The EHR biochemistry data was generated in accredited NHS laboratories for clinical use, therefore the measures are accurate, with internal and external quality control and quality assurance processes in place for all tests and investigations (SCI-Store, www.sci.scot.nhs.uk/products/store/storemain.htm). Efforts have also been made to map legacy local test codes from clinical biochemistry laboratories to the Read Clinical Classification (Read Codes)^13^. Over time assay methods, instrumentation and automation protocols will have changed, but outputs have to show consistency for clinical diagnostic purposes. An NHS Data Quality Assurance team is responsible for ensuring Scottish Morbidity Record (SMR) datasets are accurate, consistent and comparable across time and between sources (www.isdscotland.org/Products-and-Services/Data-Quality/).

The data available in the EHR in Scotland is extensive, covering both health and social domains, e.g. the Scottish Index of Multiple Deprivation, see www.ndc.scot.nhs.uk/National-Datasets/index.asp. The scope of the GS:SFHS resource is illustrated by showing the numbers of participants with various categories of EHR data available (Figure 2). However, some gaps exist. For example, data from primary care (the Scottish Primary Care Information Resource, www.spire.scot.nhs.uk/) may become available for linkage in future. This would be particularly useful for research on conditions such as dementia where much of the health service contact is with general practitioners. Another limitation is that any features of illness that occur to participants outside NHS Scotland will not be documented. However, the availability of GS:SFHS study data that was collected at recruitment, together with the range of different types of data available longitudinally in the EHR, mean that (for example) accurate classification of cases and controls can be achieved.

**Figure 2.**
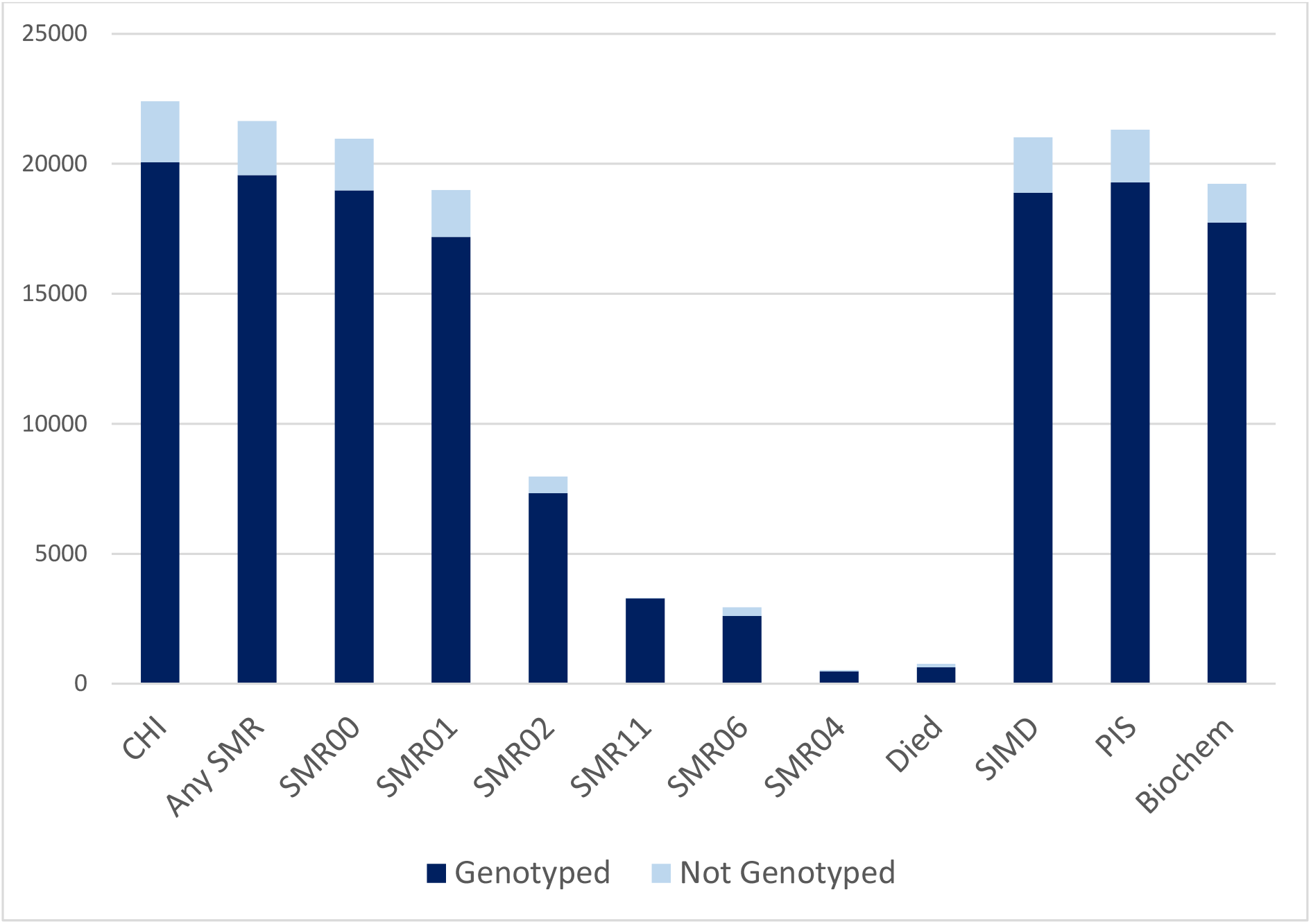
GS:SFHS participants with categories of EHR data available through CHI linkage. The light blue bars indicate the numbers of participants with linked data but not genetic data available, exact numbers given in parentheses below. The dark blue bars indicate the numbers with genome-wide genotype data (Illumina OmniExpressExome array) available in each EHR dataset. CHI, Community Health Index (n = 22,408); Any SMR, Scottish Morbidity Record (n = 21,651); SMR00, Outpatient (n = 20,777); SMR01, General acute/Inpatient (n = 18,686); SMR02, Maternity Inpatient (n = 7,804); SMR11, Neonatal Inpatient (n = 3,284); SMR06, Scottish Cancer Registry (n = 2,562); SMR04, Mental Health Inpatient (n = 498); Died (n = 768); SIMD, Scottish Index of Multiple Deprivation (n = 21,021); PIS, Prescribing Information System (n = 21,029); Biochem, biochemistry laboratory data (n = 19,233).

An illustration of how the genetic and EHR data in GS:SFHS can be used is in examining the psychiatric history of cases of major depressive disorder (MDD) and controls using record linkage to the Scottish Morbidity Record (Outpatient and Mental Health Inpatient datasets) and prescription data (for history of antidepressants). This information has been used in haplotype association analyses of MDD^14^, stratification of MDD into genetic subgroups^15^ and genome-wide meta-analyses of stratified depression.^16^ Data science has great potential as a catalyst for improved mental health recognition, understanding, support and outcomes.^17,18^ Validation of the laboratory EHR data was provided by a genome-wide association study (GWAS) of serum urate in the Tayside regional subset of the cohort.^5^ Uric acid is a medically relevant phenotype measure, with high levels leading to the formation of monosodium urate crystals that can cause gout. Hyperuricaemia has additionally been associated with a variety of diseases including type 2 diabetes, hypertension and cardiovascular disease, while hypouricaemia has been linked to neurodegenerative disorders including Parkinson’s disease and Alzheimer’s disease.^19^ GWAS of uric acid was performed using EHR-derived measures from 2,077 individuals^5^ and shows the strong signal in the *SLC2A9* gene previously reported using data gathered specifically for research^20,21,22^. This positive control confirmed that the EHR-derived biochemistry data can be suitable for population-based analyses, despite being collected for clinical purposes. The initial GWAS has now been extended into the Glasgow regional subset of the cohort (Figure 1), increasing the number of individuals with both genotype data and at least one uric acid measurement to 3,160, within the total of 19,233 participants where biochemistry data can be accessed (Figure 2). Summary statistics for ∼600,000 single nucleotide polymorphisms are available at http://dx.doi.org/10.7488/ds/2125 and these results will be included in a genome-wide association meta-analysis by the Global Urate Genetics Consortium.

Table 1 lists the 30 most frequently collected serum biochemistry measures, with the number of unique participants and total number of their tests shown. The test with the highest number of measures recorded (in unique participants) is creatinine, with 206,498 records from 17,393 participants. Urate is relatively low down the list (27^th^ position), but nonetheless enough data is available to yield a highly significant (top hit p-value = 7.29 × 10^-21^) GWAS result (Figure 1). This helps to demonstrate the breadth and depth of research that is possible with the biochemistry dataset, only one of many that are available (Figure 2).

**Table 1.**
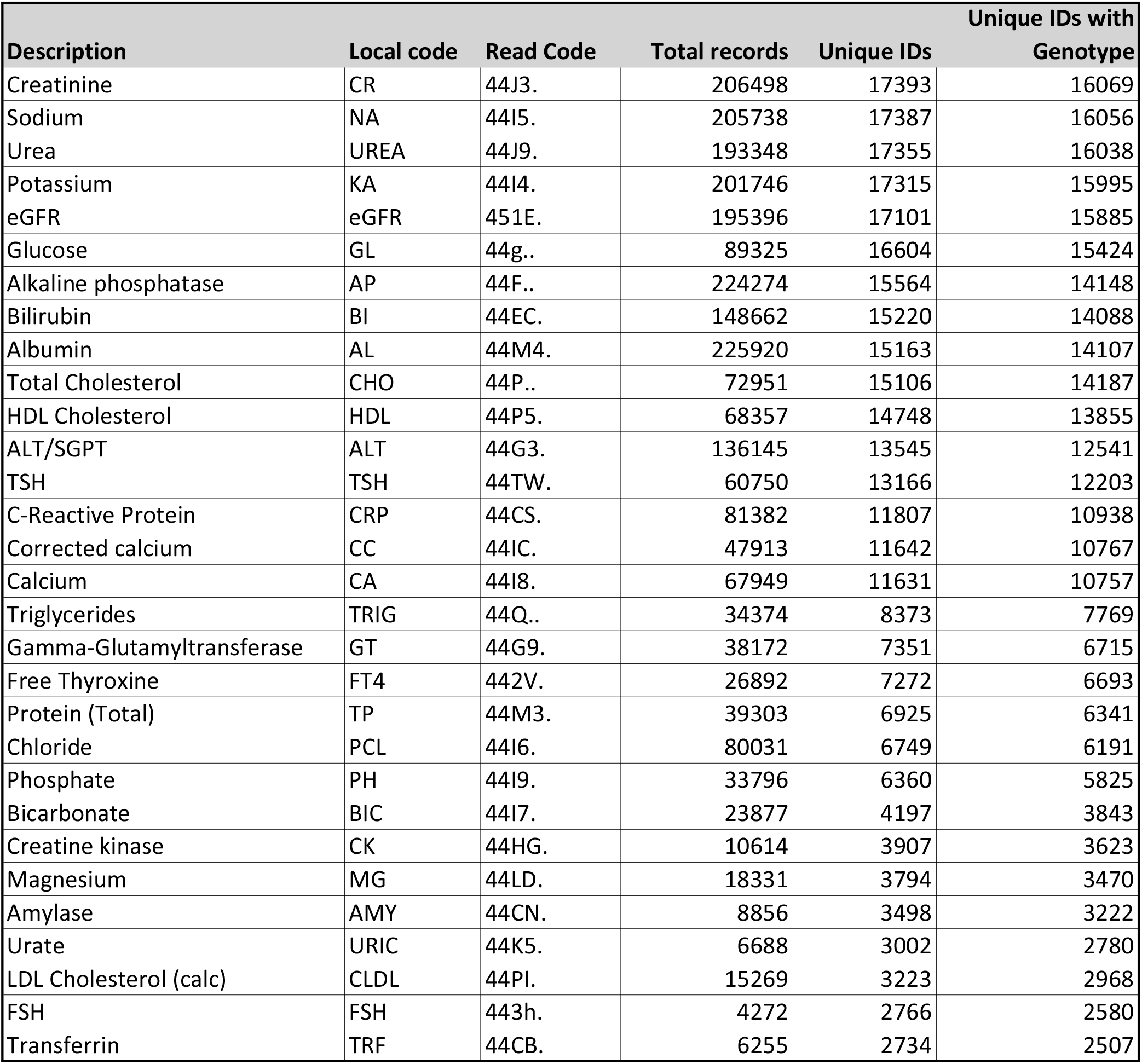
The 30 most frequently collected serum biochemistry measures, by number of unique participants. The description of each measure, local code, Read Code, total number of records, number of unique participants and unique participants with genotype data are shown. Totals are for data collected from 2006 to 2016 on participants aged 18 or over.

Hospital admission, prescription and biochemistry EHRs can be used to infer disease status in individuals after recruitment to the study has concluded, including for diseases that were not part of the initial data collection. One such example is gout, which was not explicitly included in the pre-clinical questionnaire, but with access to EHR data it is possible to ascertain gout status for participants in GS:SFHS. Not all individuals with hyperuricaemia develop gout, which may mean that other factors predispose individuals to a greater or lesser risk of disease progression. Identifying these factors could help inform targeted prevention and personalised management of the disease. The up-to-date status of risk factors available in GS:SFHS from EHR linkage makes it an excellent resource for a case-to-high-risk-control GWAS. A static study might incorrectly class an individual as a high-risk control simply because they did not develop gout until after data collection had concluded. Here, 420 gout cases have been identified through the use of urate lowering medication, obtained from the Scottish National Prescribing Information System (PIS).^23^ Additional information can be obtained from the GS:SFHS baseline phenotype dataset, including self-reported use of medications and measures such as body mass index. This information, together with the range of risk factors available in the biochemistry, prescribing and morbidity EHR datasets (e.g. gout ICD-10 codes in SMR01), will be used to select risk-matched individuals who have not developed gout for a case-control GWAS of GS:SFHS participants.

## Ethics Policies

GS:SFHS has Research Tissue Bank status from the East of Scotland Research Ethics Service (REC Reference Number: 15/ES/0040). This provides a favourable opinion for a wide range of data and sample uses within medical research. Research that includes access to individual-level EHR data is notified to the Research Ethics Committee by the GS management team on behalf of the researchers, through a notice of substantial amendment.

## Data Availability

The study phenotype (cohort profile)^1^ and genotype data (both directly typed and imputed to the Haplotype Reference Consortium release 1.1 panel)^5^ have both been described. A phenotype data dictionary is available (dx.doi.org/10.7488/ds/2057) and open access GWAS summary statistics can be downloaded (http://datashare.is.ed.ac.uk/handle/10283/2789). Non-identifiable information from the GS:SFHS cohort is available to researchers in the UK and to international collaborators through application to the GS Access Committee. Generation Scotland operates a managed data access process including an online application form (www.ed.ac.uk/generation-scotland/using-resources/access-to-resources) and proposals are reviewed by the GS Access Committee. Summary information to help researchers assess the feasibility and statistical power of a proposed project is available on request by contacting. For example, the numbers of participants with each biochemistry measure listed, and the total number of measures available per participant, are provided in Table 1. Researchers requesting individual-level EHR data must also submit their proposal to the NHS Public Benefit and Privacy Panel for Health and Social Care (www.informationgovernance.scot.nhs.uk/pbpphsc/home/for-applicants/).

If access to biochemistry EHR data is part of the proposed research, an additional application to the two data Safe Havens holding this data is required (www.nhsggc.org.uk/about-us/professional-support-sites/nhsggc-safe-haven/ and www.dundee.ac.uk/hic/hicsafehaven/), with a new safe haven workspace created for each project. The biochemistry data was generated as part of routine medical care by the NHS, and only modest cost recovery charges are made to provide access to it for research purposes. This compares favourably with the costs of commissioning new tests on blood, serum or plasma samples. The Generation Scotland data access process also incurs an administrative cost recovery charge. Once a proposal has been approved, researchers are provided with pseudo-anonymised data extracts.

CHI numbers are replaced by unique study numbers and personal identifying information is removed. Numbers of participant records currently available for each of the main EHR data categories, and the proportion with genome-wide genotype data, are illustrated in Figure 2. The numbers of data points will increase over time as the participants grow older, covering a wide range of clinically relevant outcomes and continuing to extend this rich research resource.

## Consent

Only data from those GS:SFHS participants who gave written informed consent for record linkage of their GS:SFHS study data to their medical records are used.

## Acknowledgements

We are grateful to all the families who took part in GS:SFHS, the general practitioners and the Scottish School of Primary Care for their help in recruiting them, and the entire participant recruitment team. We thank staff at the University of Dundee Health Informatics Centre, NHS Greater Glasgow and Clyde Safe Haven and NHS National Service Scotland eDRIS for their expert assistance with EHR data linkage. The work in this paper uses data provided by patients and collected by the NHS as part of their care and support.

